# Quantifying the wave resistance of a swimmer

**DOI:** 10.1101/2020.06.22.164236

**Authors:** Thomas A. J. Dickson, Dominic Taunton, Joe Banks, Dominic Hudson, Stephen Turnock

## Abstract

Quantifying the wave resistance of a swimmer as a function of depth assists in identifying the optimum depth for the glide phases of competition. Previous experiments have inferred how immersed depth influences the drag acting on a swimmer [1], but have not directly quantified the magnitude of wave resistance. This research experimentally validates the use of thin-ship theory for quantifying the wave resistance of a realistic swimmer geometry. The drag and wave pattern of a female swimmer mannequin were experimentally measured over a range of depths from 0.05m to 1.00m at a speed of 2.50 m/s. Numerical simulations agree with experiment to confirm that there were negligible reductions in wave resistance below a depth of 0.40m. Larger swimming pool dimensions are shown to be significant at reducing wave resistance at speeds above 2.0 m/s and depths below 0.40m. Truncating the swimmer’s body at the upper thigh increases the wave resistance at speeds below 2.0m/s but is not significant at higher speeds, indicating that the upper body is the main contributor to the wave system. Numerical experiments indicate that rotating the shoulders towards the surface is more influential than the feet, demonstrating the impact of the upper body on wave resistance.

## 1 Introduction

As swimmers spend 10 − 25% of their time in the glide phase in competition [2] reducing the drag experienced in the glide phase will improve competitiveness. One method of reducing drag is by improving technique through reducing the drag associated with the generated wave pattern, also known as the wave resistance. Wave resistance has been identified as a key source of resistance in the glide phase [3]. Figure 1 shows an athlete in the glide phase. The influence of wave resistance on total swimmer drag has been inferred by measuring changes in the drag of a swimmer mannequin as its immersion increases [1], but has not been directly experimentally quantified. Linear slender body theory, also known as thin-ship theory, can be used to predict the wave resistance of ships [4] but has not been validated against swimmer geometries. After validating the use of linear slender body theory on swimmer geometries it will be possible to use it to directly quantify swimmer wave resistance.

**Figure 1:**
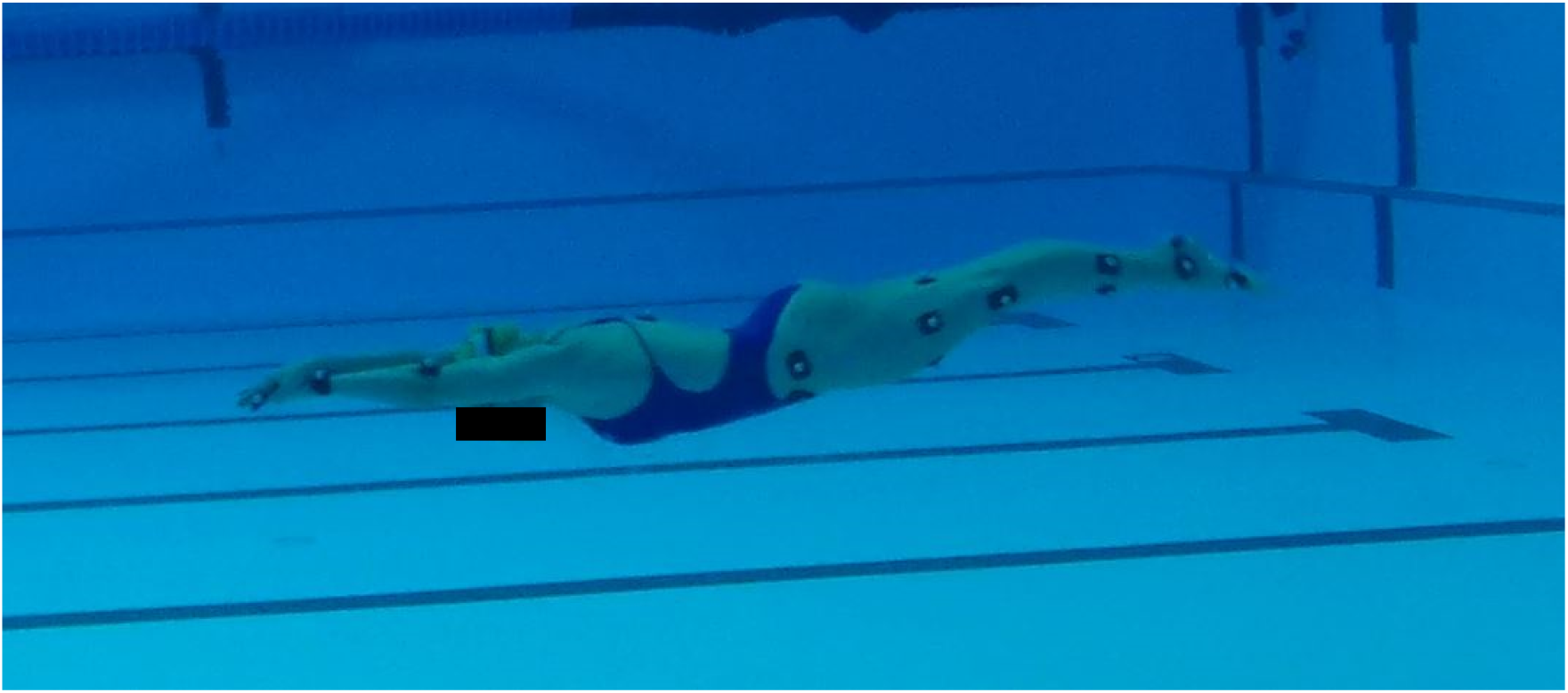
Female athlete in the glide phase.

Two key mechanisms contribute to the resistance of a swimmer: the viscous resistance, *R_v_*, which is Reynolds number dependent and the wave resistance, *R_w_*, which is Froude number dependent [5]. The Froude number is defined as 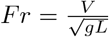, where *g* is the gravitational constant, *L* is the waterline length and *V* is the speed. The Reynolds number can be defined as 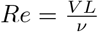, where *ν* is the kinematic viscosity of fluid. Figure 2 illustrates the breakdown of different sources of hydrodynamic resistance and how they can be analysed.

**Figure 2:**
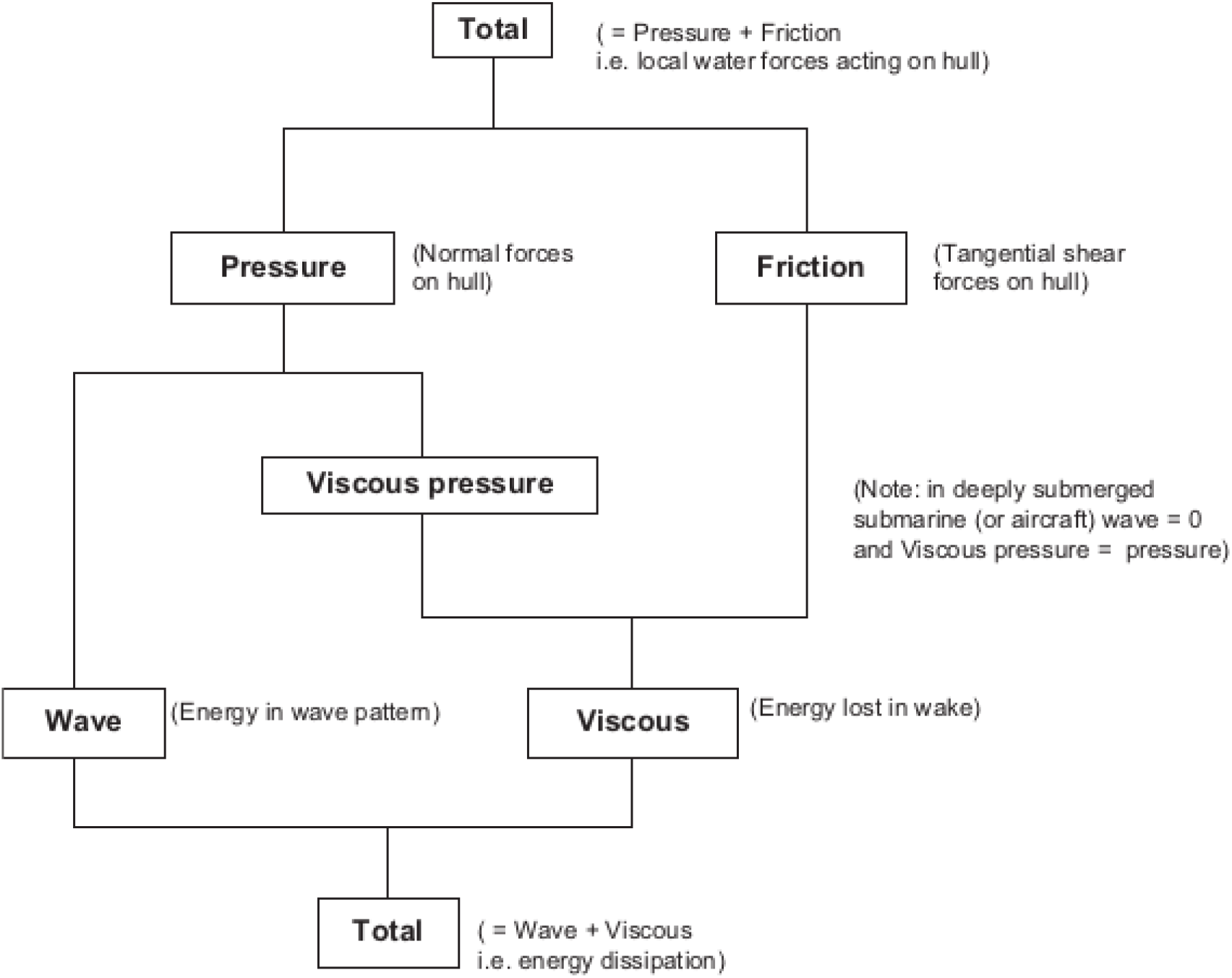
Breakdown of the physical components of the resistance of swimmers [5, p. 14]. This illustrates how the total resistance of a swimmer can be decomposed into wave and viscous resistance.

Different experimental and numerical methods may be used to measure the wave resistance of a swimmer. Computational Fluid Dynamics (CFD) has been used to model the total resistance of the swimmer on the surface, [6, 7]. However, the individual resistance components of a swimmer have yet to be experimentally validated. Thin-ship theory has been successfully applied in conventional naval architecture for quantifying the wave resistance of ships [8, 9], for example, but has yet to be applied to swimmer geometries. Thin-ship theory offers the opportunity for accurate results with run times that are two to three orders of magnitude faster than CFD [5].

This paper presents the results of an experimental and numerical investigation into quantifying the influence of swimmer depth, trim and physique on the magnitude on the wave resistance. Experimental measurements of the drag and wave pattern of a swimmer mannequin at different depths are used to validate the application of thin-ship theory to swimmer geometries. Thin-ship theory is then used to model the influence of trim, swimmer physique and swimming pool dimensions. The wave patterns for athletes in the glide phase are predicted and key depths are identified for optimising performance.

## 2 Thin-ship theory

Thin-ship Theory (TST) has been used to estimate the wave resistance of ships for several decades [8]. Development to initial work [10, 11, 12] have enabled the application of TST to vessels of a variety of shapes. Some examples of applying TST to ships include [8, 9]. As TST has been shown to provide fast accurate simulations of ship hydrodynamics, applications of TST to swimmer hydrodynamics have also been explored.

TST has been used to quantify the wave resistance of a male swimmer being towed at speeds of 1.7m/s and 2.1m/s [13]. Experimental estimates of wave resistance were obtained from wave cuts using the matrix method [8]. Despite good agreement between numerical and experimental results, there were areas of significant experimental and numerical uncertainty. A source of experimental uncertainty was the inability of the human athlete to maintain a constant shape throughout each run. Furthermore, the influence of the pools infinity edge was not modelled using CFD, contributing towards the uncertainty in the numerical predictions.

A more comprehensive numerical study investigated the wave resistance of “swimmer-like forms” travelling singly and in groups [14]. This study primarily focused on the drag reductions experienced by swimmer like forms moving in wake regions of other swimmer like forms. No experimental verification of these results were provided and the swimmer-like forms did not have realistic human geometries.

Several assumptions must be met in order to model the resistance of a swimmer using TST. The swimmer is assumed to be slender, have a high length to breadth ratio, the fluid is inviscid, incompressible and homogeneous, the fluid motion is steady and irrotational, the surface tension of the fluid may be neglected and the wave height at the free surface is small compared with the wave length. In this context, the assumption of high length to breadth ratio assumes that the body of the athlete being modelled is slender.

In thin-ship theory, bodies are represented by planar arrays of Kelvin sources on the local centre line in a channel of finite breadth and depth [5]. The body shape is represented by mapping a series of triangular or quadrilateral panels onto its surface. The strength of the source on each panel is calculated from the panel normal, Equation 1. 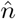 is the outward unit normal vector of the panel and the constant onset swimming speed *U* = (−*U_x_*, 0, 0,), with the x-axis pointing from toe to head, z vertically downwards and is fixed with the body. The panelled body is generated from a scan of a prone athlete.

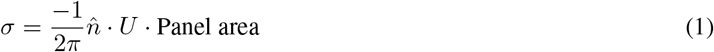

The wave resistance of the sources is described in terms of far-field Eggers coefficients for a source in a finite channel [12] using an expression derived by [8]. Equation 2 calculates the total wave resistance which is dependent on the wave harmonic, *m*, where the wave elevation for a given harmonic *ζ_m_* is given by Equation 3.

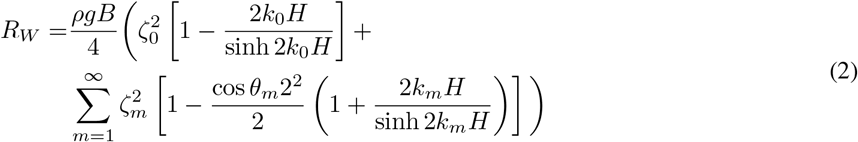

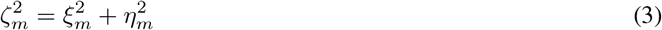

The elevation terms for a source at (*x_σ_*, *y_σ_*, *z_σ_*) are given in Equation 4. Note that the term for *m* = 0 is halved and that the last cosine term applies to even *m* and the sine term to odd *m*. The summation represents the effect of all the sources describing the hull, typically 800 [9].

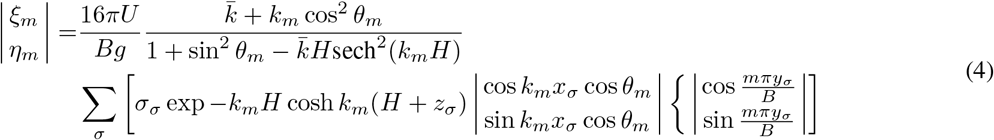

The fundamental wave number is given by 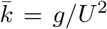. The wave number *k_m_* and the wave angle Θ_*m*_ of the *m*th harmonic satisfy the wave speed condition and wall reflection condition, described in Equations 5 and 6, respectively.

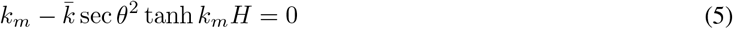

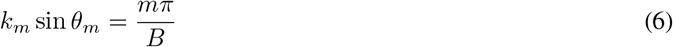

Thin-ship theory can be used to provide an estimate of the proportion of transverse and divergent content in the wave system which can be compared with values derived from physical measurements of the experimental wave elevation. It is possible to relate the wave pattern and associated drag with the length Froude number. A parameter used to describe the influence of shallow depth on resistance is the depth Froude number, 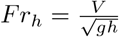, where *h* is the depth underneath the ship or swimmer [5, p. 97]. At *Fr_h_* < 1.0 the wave system has a transverse wave system and a divergent wave system propagating away from the ship at an angle approximately 35°. Figure 3 shows how the wave system of a moving ship can be decomposed into transverse and divergent wave systems. As the speed approaches the critical speed, *Fr_h_* = 1.00, the divergent wave angle approaches 0°. As gravity waves cannot travel faster than 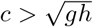 they are left behind at speeds *Fr_h_* > 1.00 leaving a divergent wave system.

**Figure 3:**
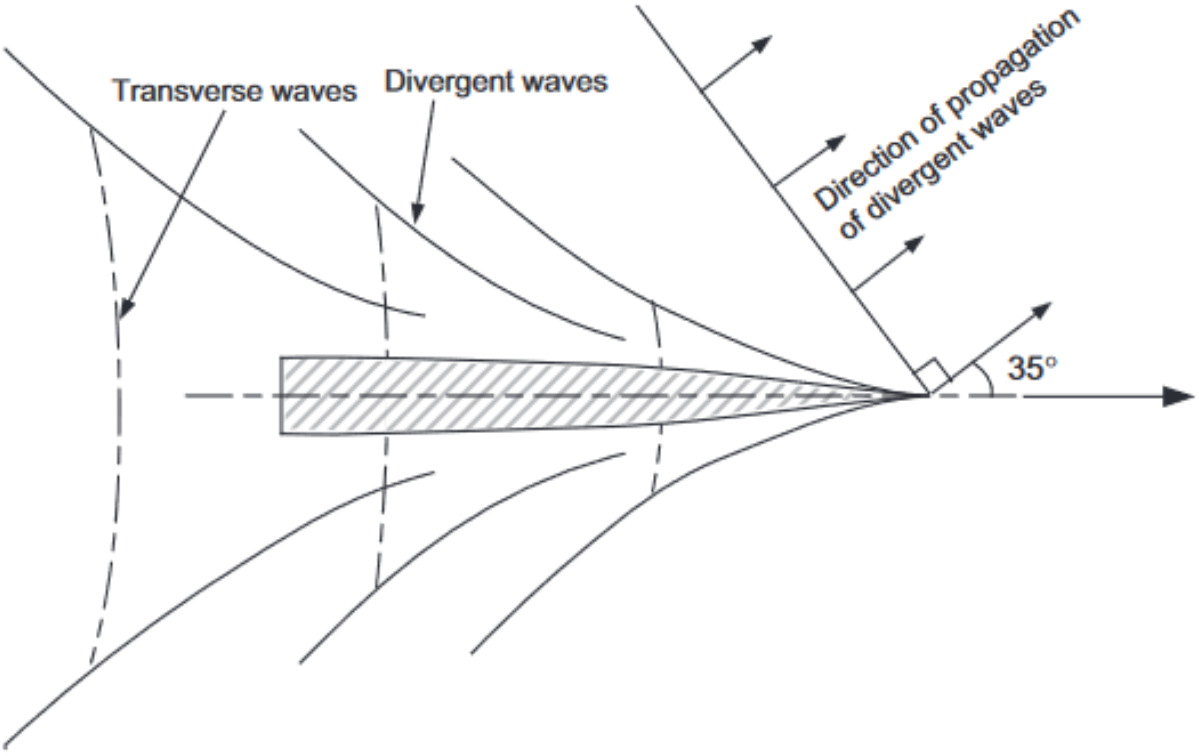
Divergent and transverse wave systems for a ship moving at a constant *Fn_h_* [5]. These wave systems are generated for any object moving along the water and air interface.

### 2.1 Passive resistance of swimmers

A large body of research has focused on quantifying the passive drag of a swimmer [2]. However, there has been less focus on quantifying the magnitude of wave resistance which contributes to the total passive drag that acts on a swimmer: 1. experimental techniques using a flume or towing swimmers or mannequins [15, 1, 13], 2. Numerical simulations [16], and 3. analytical methods drawing from naval architectural techniques [17, 13, 14]. Of specific interest is how these methods are employed to investigate the influence of wave resistance on the passive drag of a swimmer.

Initial research quantified that it was possible to travel 1.93 m/s after pushoff at the surface compared to 2.24m/s at a depth of 0.6m, a 16% reduction in speed [15]. This experiment illustrated how increasing depth reduces the total resistance of the swimmer. A further study inferred that wave drag was a significant component of total drag of an athlete [3], although this study assumed that wave drag was negligible below 1.6 m/s and only recorded the force generated from each arm stroke. The drag acting on a mannequin in a flume was measured for a range of depths and speeds [1] and identified that the total passive drag experienced no significant reduction below a depth of 0.75 m. Although comprehensive, this study did not directly quantify the wave resistance component.

More recent research simulated the drag on a swimmer at a range of speeds and depths using CFD and also found that there was no significant reduction of drag beyond a depth of 0.75 m [18]. However, there was no comparison with experimental results or verification of the wave pattern generated. The application of computational fluid dynamics to modelling swimmers is limited by the difficulties using CFD to model the same boundary conditions seen in the swimming pool, for example, the infinity pool edge [2].

The wave resistance of an object may be estimated based on a recording of the height of the wave pattern the object generates as it moves at a constant speed. This is known as taking a “wave cut”, and multiple “wave cuts” are taken at different transverse locations to estimate the energy required to generate the wave pattern [5].

Thin-ship theory can be used to estimate the wave resistance of a hull form [5, p. 177] and has been well applied in conventional naval architecture [9]. The wave resistance of other immersed objects, such as submarines, has also been investigated, [19] for example. It has been found that the diameter, length and depth all influence the wave making resistance. Reported experiments are usually conducted at lower Froude numbers than those seen in competitive swimming, indicating that new experiments are required to give results with meaningful context. In this paper we investigate the influence of swimmer depth on the wave resistance as a function of depth.

## 3 Experimental Method

### 3.1 Mannequin

A mannequin was constructed of an elite female athlete in the prone position for hydrodynamic testing, from a 3D scan. The same scan was converted into a triangulated mesh model. The lower half of the legs were removed from the model in order to provide mounting points for the support structure, known as “stings”. One of the challenges associated with experimentally quantifying the forces acting on a submerged object is the need for a physical connection which can affect the flow. Any connection upstream will necessarily change the flow over the tested object therefore it was decided to place the support structure down stream of the swimmer geometry. The manufactured mannequin geometry is henceforth referred to as the “truncated swimmer geometry”.

The truncated swimmer geometry was 3D printed and a finish of resin paint was applied. Multiple versions of the head and arms were manufactured and trialled. Gaps between the arms, heads and torso were covered using foil tape to ensure a smooth surface. Two foils were manufactured and placed on the stings at the interface between fluid and air to reduce in drag. The particulars of the mannequin and the foils used to moderate the flow past the support structure are described in Table 1. An initial study identified that using more than 4184 panels to represent the mannequins geometry in thin ship theory had a negligible influence on the simulation results.

**Table 1:** Particulars of the foils and mannequin used.

| Parameter | Foil | Truncated Mannequin | Full Mannequin |
| --- | --- | --- | --- |
| Length (m) | 0.3 | 1.58 | 2.3231 |
| Breadth (m) | 0.0399 | 0.39 | 0.3944 |
| Draught (m) | 0.25 | 0.30 | 0.3022 |
| Centreline | 0.075 | 0.00 | 0.0 |
| Wetted Area (m<sup>2</sup>) | 0.1517 | 1.41 | 1.8265 |
| # Panels for TST | 400 | 4184 | 5444 |

Measurements were taken at various points over the truncated mannequin at a *D* = 0.05m datum, Point C in Figure 4, where *D* is the depth of immersion. Table 2 records the depth of immersion at 5 locations on the mannequin and support structure. The physical locations of the measurement points are marked on Figure 4.

**Table 2:** Truncated mannequin free surface measurements from 0.05m test height.

| Location | Measurement (m) | Diagram |
| --- | --- | --- |
| Right middle finger | 0.17 | A |
| Back of the head | 0.04 | B |
| Mark on the back of shoulder blades | 0.05 | C (Datum) |
| Stern mark (between legs) | 0.125 | D |
| Rear right box section top | 0.15 | E |

**Figure 4:**
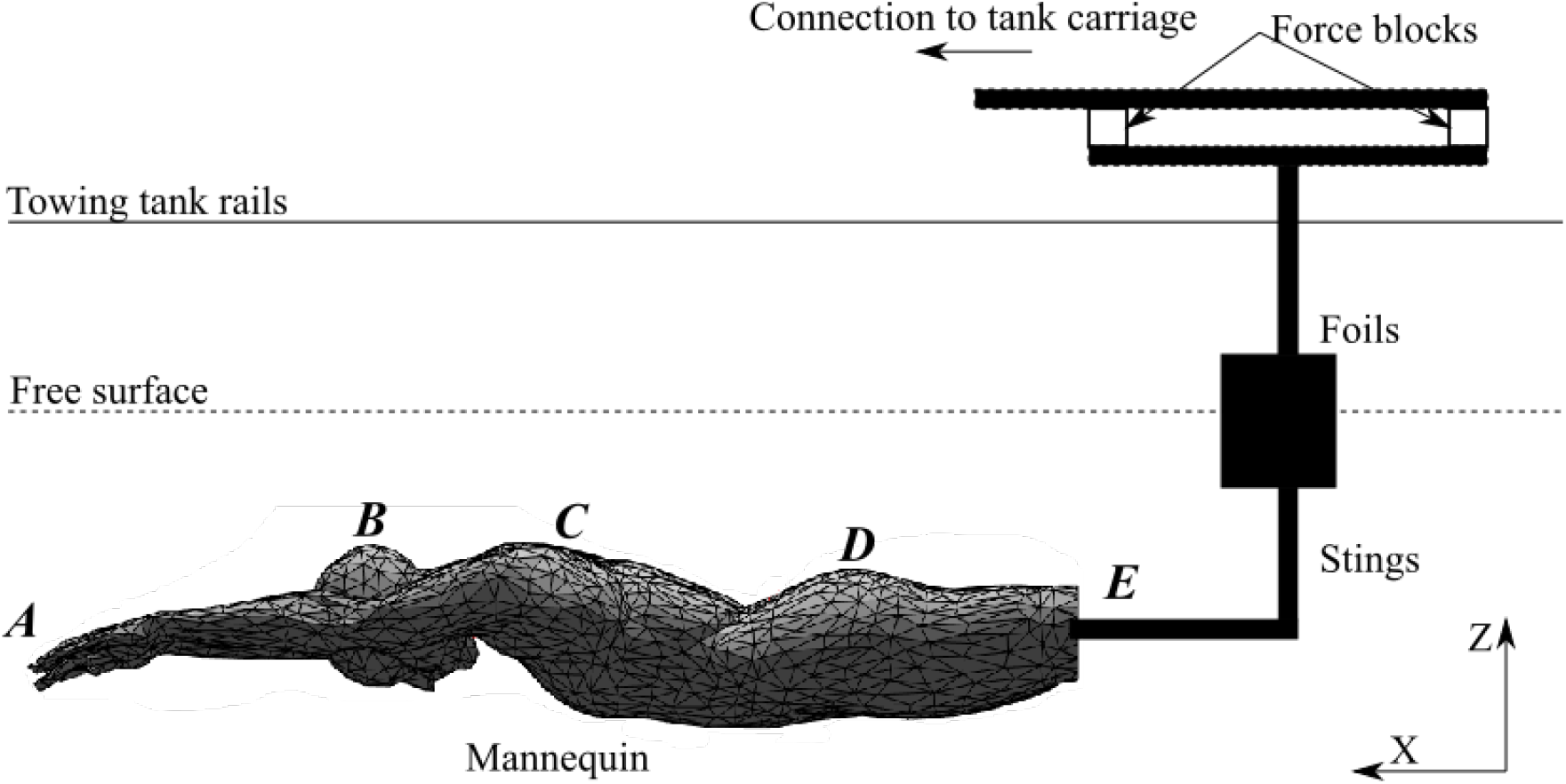
Diagram of the experimental set up in the X-Z plane.

### 3.2 Force and wave set measurement

The experiments were carried out at the Southampton Solent towing tank at a temperature of 15°, which has the particulars described in Table 3. The mannequin is positioned beneath the towing tank carriage in the manner described in Figure 4. The mannequin is fixed to two stings which pierce the free surface. A foil is fixed at the interface of the free surface to reduce the wave resistance of the structure and make it easier to model in TST. A wave probe array is used to record the wave pattern generated in each experiment. The wave probes were placed 1.11m, 1.207m and 1.291m from the centreline of the tank.

**Table 3:**
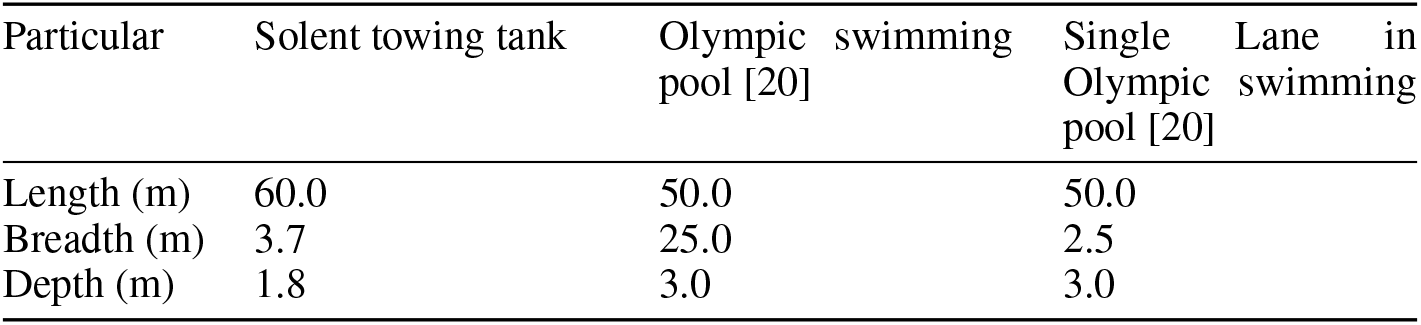
Particulars of the different channels used in experiment and modelling.

| Particular | Solent towing tank | Olympic swimming pool [20] | Single Lane in Olympic swimming pool [20] |
| --- | --- | --- | --- |
| Length (m) | 60.0 | 50.0 | 50.0 |
| Breadth (m) | 3.7 | 25.0 | 2.5 |
| Depth (m) | 1.8 | 3.0 | 3.0 |

**Table 4:** Variation in calibration factors for quantifying experimental uncertainty.

| Variation<br>(mm/V) | WP 1 | WP 2 | WP 3 |
| --- | --- | --- | --- |
| -1 | -51.707 | -48.838 | -45.729 |
| -0.1 | -50.707 | -47.838 | -44.729 |
| 0 | -50.607 | -47.738 | -44.629 |
| 0.1 | -50.507 | -47.638 | -44.529 |
| 1 | -49.607 | -46.738 | -43.629 |

The support structure can alter the depth of immersion and trim of the mannequin. The drag force acting on the mannequin and support structure was measured using four flexure based force blocks fitted with displacement transducers. The force transducers were energised and conditioned using RDP 621 analogue cards. A low pass filter was applied to the data, which was recorded at 100 Hz using a custom MATLAB program. The scantlings of the support structure for the stings positioning the mannequin were sized to ensure the deflection of the structure is less than 1mm for the ranges of forces it would experience in testing.

Three longitudinal wave cuts were recorded during each run using an array of resistance-type wave probes mounted on the side of the towing tank. The voltage of the wave probes was recorded by a HR Wallingford wave monitor which has an inbuilt 20 Hz low pass filter. The wave probes were calibrated by mapping the change in voltage to a known change in water level vertical displacement. The wave pattern resistance was estimated using the matrix method of wave analysis from the wave cuts recorded [8].

## 4 Thin-Ship Theory for Swimmers

The drag forces and wave system of a swimmer mannequin were recorded whilst the swimmer mannequin was towed at a constant speed. A speed of 2.5m/s was chosen as it is representative of a typical glide speed seen in competition. The resistance of the truncated mannequin and the stings over a series of depths at a constant speed of 2.5m/s is shown in Figure 5.

**Figure 5:**
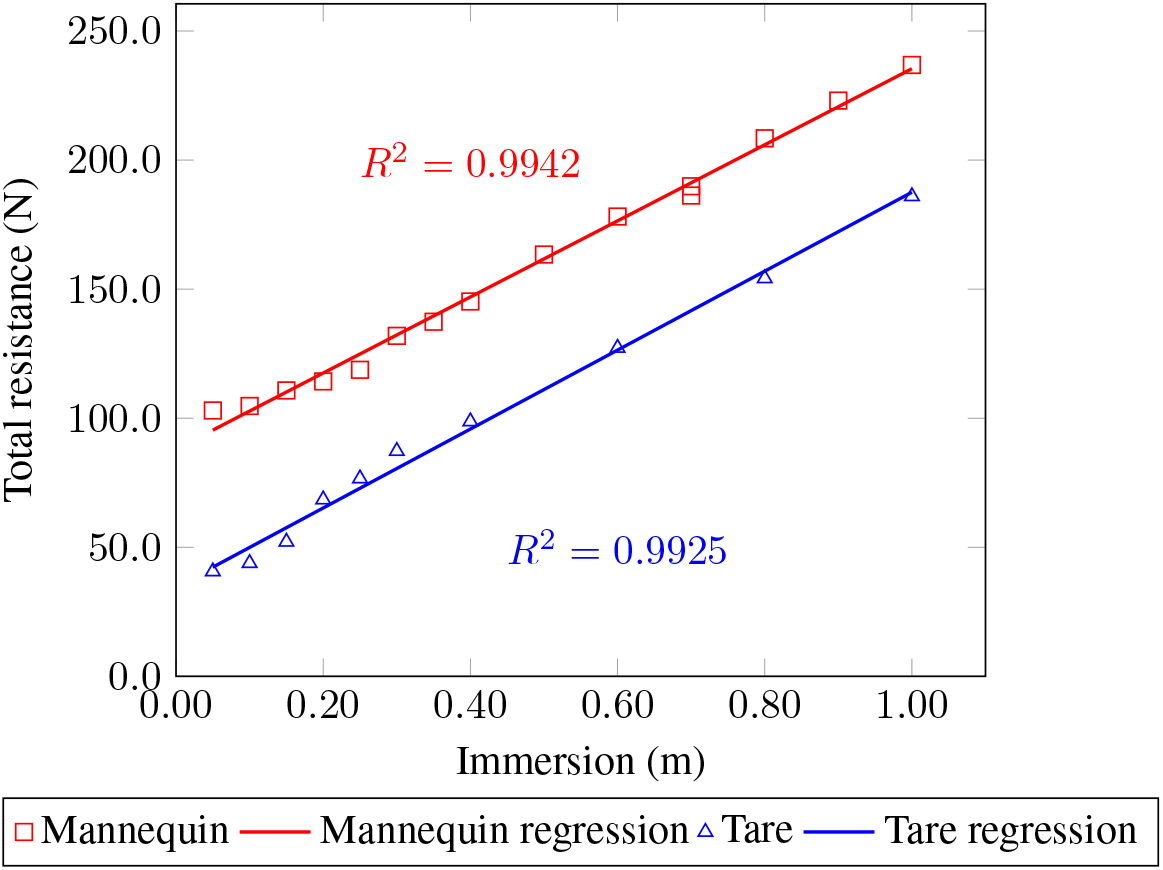
The total resistance, *R_T_*, of the mannequin and tare configurations over a range of immersed s, D.

Figure 5 shows that the drag of the mannequin and tare configurations increased linearly with depth. The reduction in mannequin drag at the depths from 0.05m to 0.40m can be attributed to the reducing wave pattern drag due to increased immersion and the drag increase due to the increasing immersed area of the support structure with depth. The wave pattern of a swimmer geometry without support structure is investigated in Section 5.3.

The wave resistance, *R_W_*, was quantified experimentally, using two methods. The first method uses the wave trace from the single probe closest to the model. The second method uses the wave trace from all three probes, known as the matrix method [8]. It is likely that the wave probe closest to the mannequin would provide overestimates of the wave resistance due to being situated in the local pressure field, whereas the analysis in Equations 2 and 3 relies on the assumption that the probes are located in in the far field wave pattern.

The experimental and theoretical wave resistance reduces significantly across the range of depths tested, Figure 6. It can be seen that thin-ship theory over estimates the wave resistance at very shallow depths but produces similar results as the depth increases. The experimental matrix method wave resistance reduces from 16.30N at *D* = 0.05m to 4.11N at *D* = 0.40m with no significant change as the depth increases further. The theoretical wave resistance reduces from 48.31 N at *D* = 0.05m to 6.80N at *D* = 0.40m and finally to 1.49N at *D* = 1.00m. The discrepancy between experimental and theoretical wave resistance at shallower depths may be due to the generation of breaking waves which are not captured in the wave trace.

**Figure 6:**
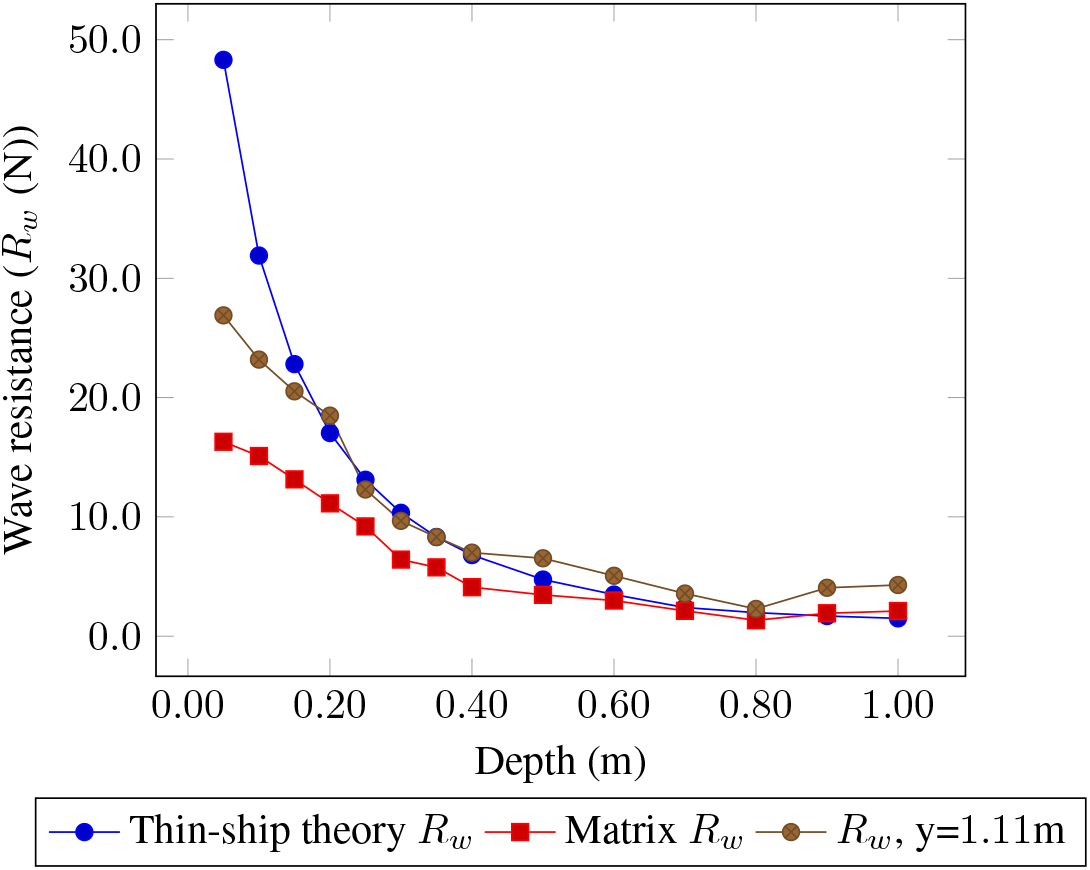
Experimental and theoretical wave resistance, *R_w_*, of the mannequin.

The wave patterns for mannequin depths of 0.05, 0.40 and 1.00 m have been calculated and they are shown in Figures 7a, 7b and 7c. There is a reduction in the angle of the divergent wave angle and the magnitude of the transverse wave between Figure 7a and Figure 7b. There is a further slight reduction in the divergent wave angle between Figure 7b and Figure 7c. Figure 7 shows that there is a significant change as the depth changes from 0.05m to 0.40m, but there is only a slight change as the depth increases further. There is a reduction in the angle of the divergent wave and an increase in the wave height. The significant change in the wave patterns shown in Figure 7 from Figures 7a and 7b to Figure 7c is most likely due to an effective change in Froude number moving from body and foils to effectively just the foils. Figure 7d shows that the foils alone generate a divergent wave pattern. However, the change in wave components which constitute each wave pattern is not known.

**Figure 7:**
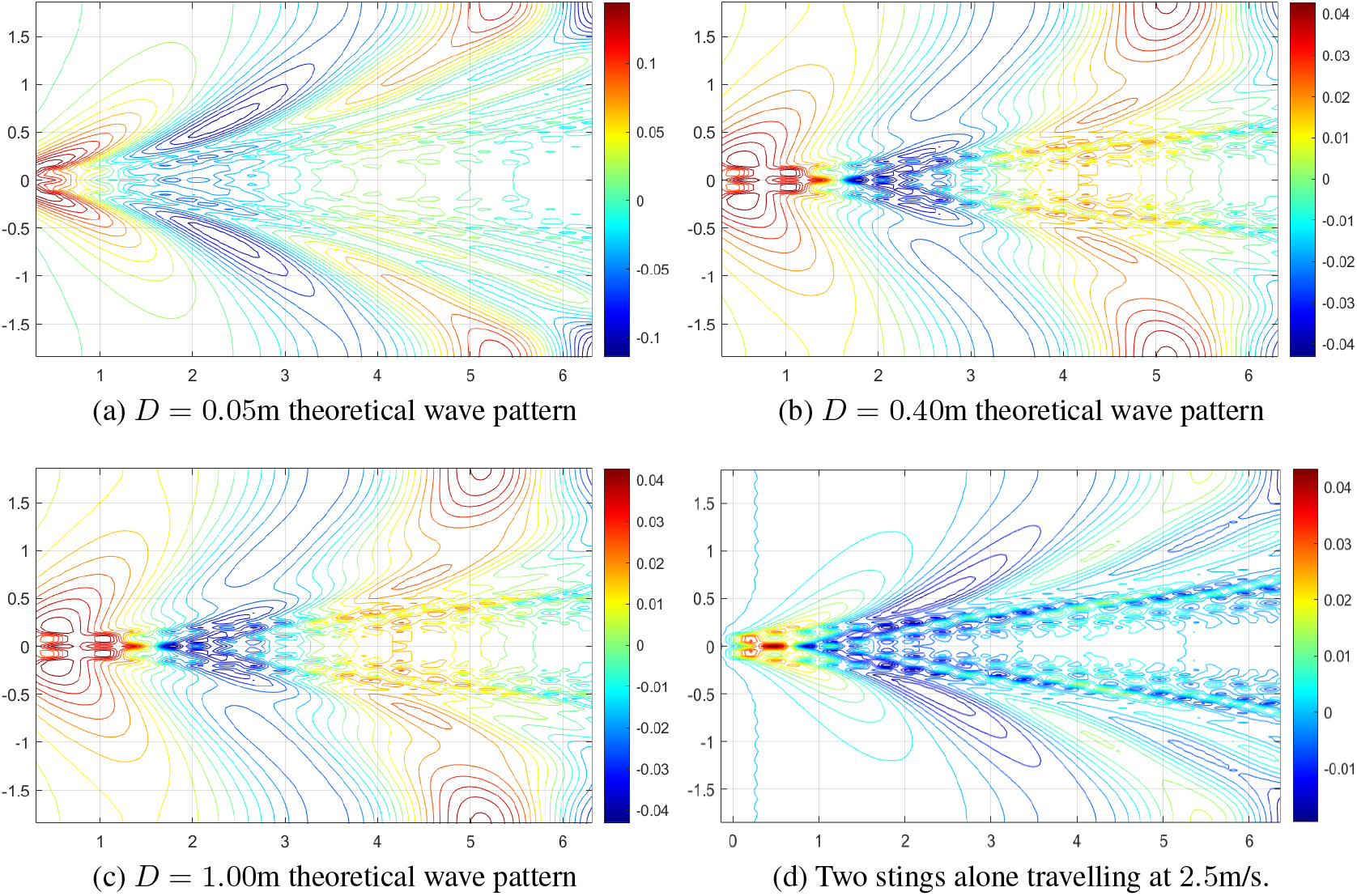
Experimental and theoretical wave patterns for the mannequin and foil combinations at depths of *D* = 0.05, 0.40 and 1.00m at a speed of 2.5m/s. The wave pattern for the foils alone is shown in Figure 7d.

The wave components recorded from the experimental and theoretical wave cuts for the mannequin and foil combination for 0.05, 0.40 and 1.00m depths are shown in Figure 8. For *D* = 0.05 the experimental wave pattern has a much high proportion of the wave system at lower wave angles, 32.5%/38.9% (theoretical/experimental) of the total wave energy for the wave component at 40°. At low depths it is possible that TST might not fully capture the wave generated at the head and shoulders. The difference between the experimental and theoretical wave components at an angle of 40° is smaller at a depth of 0.4m (17.9%/32.9%) and further reductions are seen at a depth of 1.00m (13.3%/20.9%). Note that for each depth the theoretical simulation estimates a much larger percentage of the total wave energy in the divergent wave system than is captured in reality. Wave components for all cases above 50.0° follow a similar trend to the wave components of the foil. It appears that the wave system generated by the foils dominates the high wave angle components for the mannequin and foil combination. There is a difference between the magnitude of the percentage of the wave resistance in the theoretical and experimental wave resistance for the divergent wave pattern seen at low mannequin depths. This difference is likely due to the generation of breaking waves in reality which are not generated in thin ship theory simulations.

**Figure 8:**
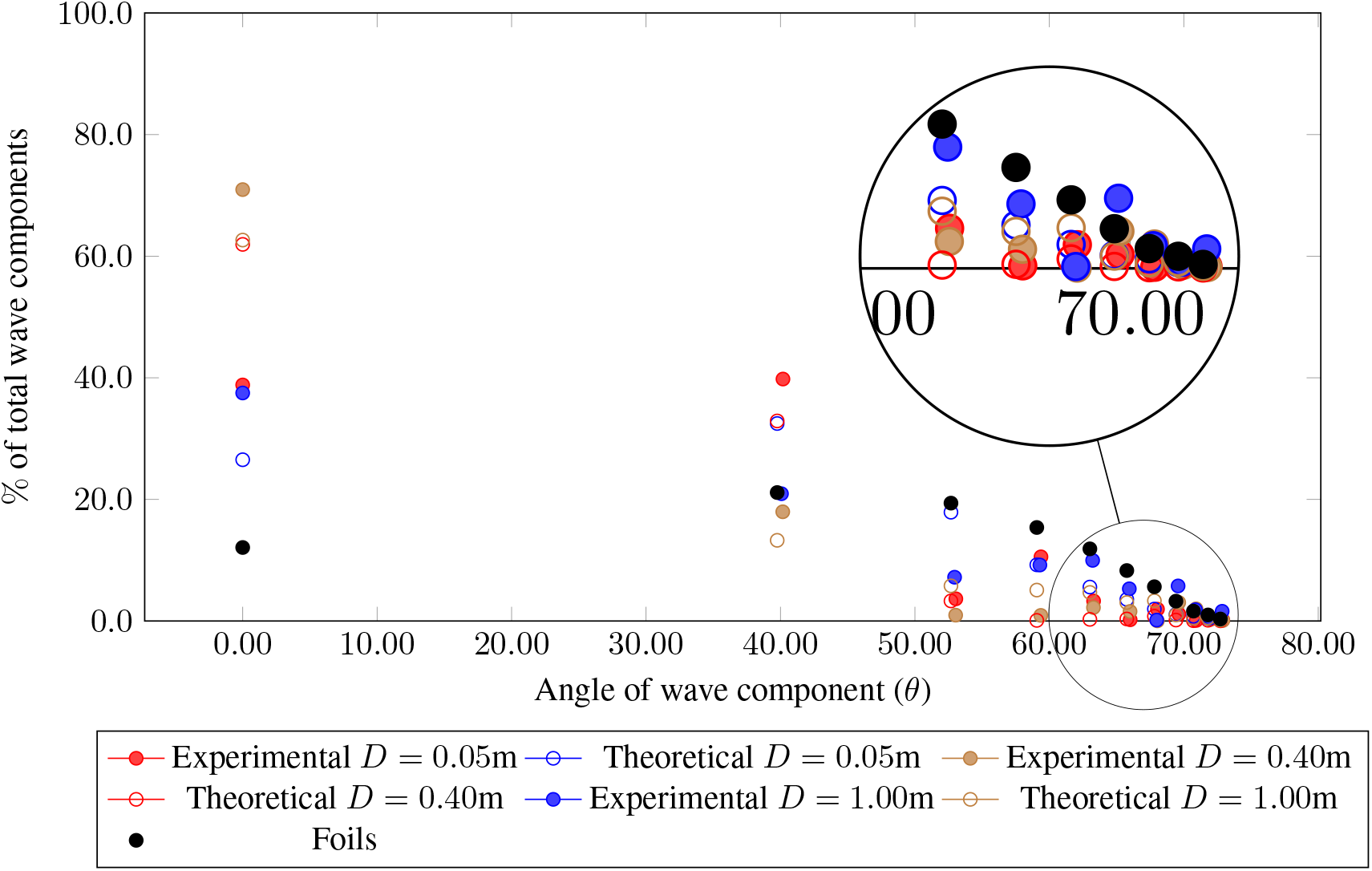
Distribution of wave resistance components for the theoretical and experimental cases at a speed of 2.5m/s.

The depth of the water is known to influence the wave system generated by the passage of an object through the water. The experiments conducted at depths of 0.05m and 1.0m are associated with depth Froude numbers of 0.610 and 0.899. *Fn_h_* values below 1.00 are subcritical, therefore shallow water effects are not considered significant for these experiments.

This section has presented experimental and theoretical results predicting the wave resistance of a swimmer mannequin and support structure combination travelling at a speed of 2.5m/s. The wave resistance is seen to reduces as the depth of the swimmer mannequin increases. The theoretical wave resistance calculated using thin ship theory is seen to under-predict the wave resistance at low depths of immersion. The small discrepancy appears to be caused by the inability to measure the wave energy lost in the breaking waves generated in the divergent wave set. TST is suitable for application to swimmer geometries as the difference between experiment and theory is small and well understood.

## 5 Parametric investigations

A series of numerical investigations have been carried out into swimmer wave resistance. Section 5.1 simulates the influence of different depths and speeds on the wave resistance produced by a truncated mannequin geometry. The impact on wave resistance of the legs is investigated by comparing the full and truncated mannequin geometry in Section 5.2. The influence of different pool dimensions on full swimmer geometry wave resistance is reported in Section 5.3, results include the predicted wave patterns of a swimmer in a swimming lane. The numerical investigations are concluded with a study of the influence of swimmer trim on the wave resistance, described in Section 5.4.

### 5.1 Influence of depth

The influence of depth and speed on truncated mannequin and sting wave resistance was quantified using TST over a range of speeds and depths. The depth was increased from 0.05m to 1.00m in increments of 0.05m and the velocity was varied between 1.00m/s to 3.00m/s in increments of 0.25m/s. Figure 9b shows how the magnitude of wave drag is larger at lower depths as the speed of the swimmer increases. Reductions of up to 20N can be achieved at speeds higher than *Fr* = 0.20 by increases in depths of 0.20m. Large reductions in wave resistance have been identified for this mannequin. Identifying how these reductions in wave resistance relate to the change in wave pattern is the focus of Section 5.3.

**Figure 9:**
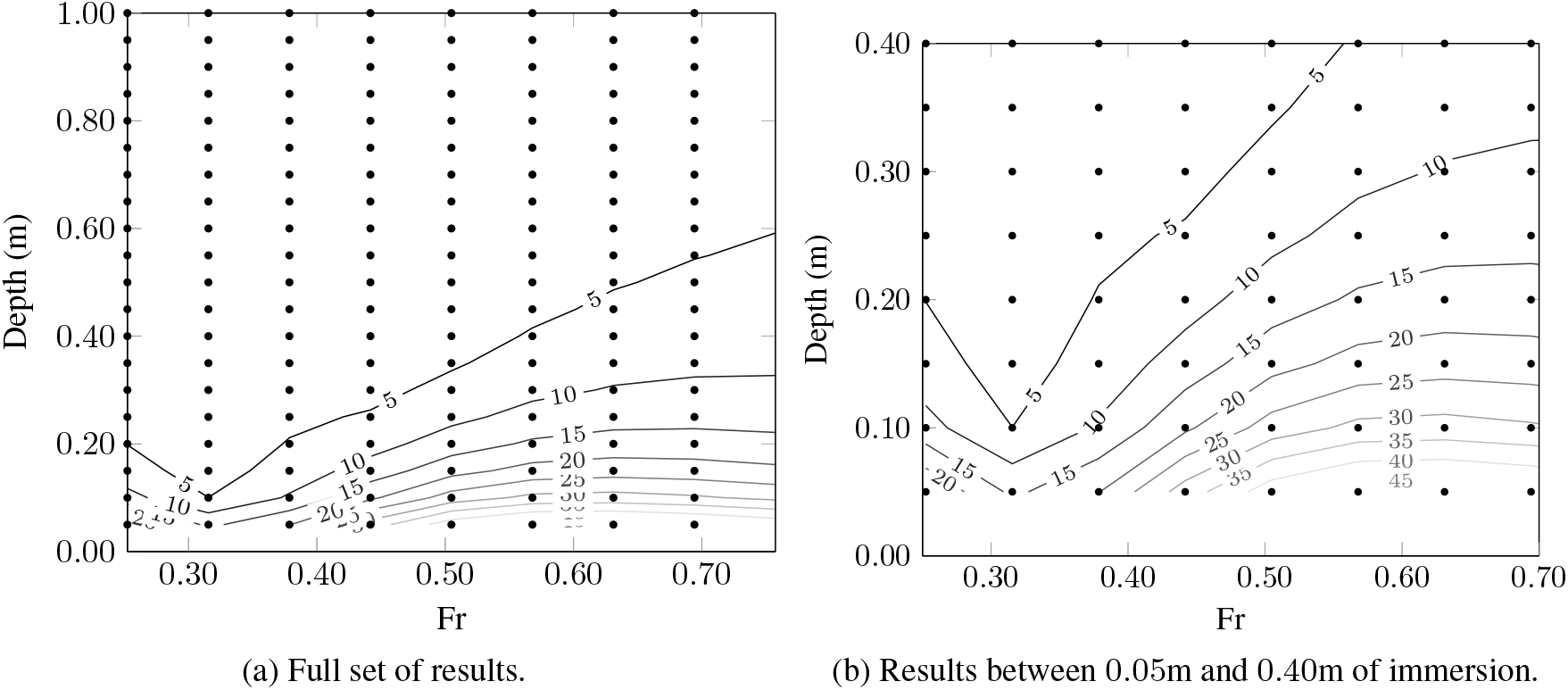
Wave resistance (N) for depths between a) 0.05m and 1.00 and b) between 0.05m to 0.40m in increments of 0.05m and speeds between 1.00m/s and 3.00m/s in increments of 0.25m/s.

### 5.2 Influence of geometry

The Froude number of an athlete is reduced by removing the legs as the waterline length is reduced. The relative increase in Froude number when investigating a full scale swimmer changes the wave pattern which causes a change in the magnitude of the wave resistance. The influence of including legs on the truncated swimmer geometry was investigated by quantifying the wave resistance of the full and truncated swimmer geometries in an open Olympic swimming pool. The dimensions of the Olympic swimming pool are recorded in Table 3 and the full geometry of the athlete is shown in Figure 10.

**Figure 10:**
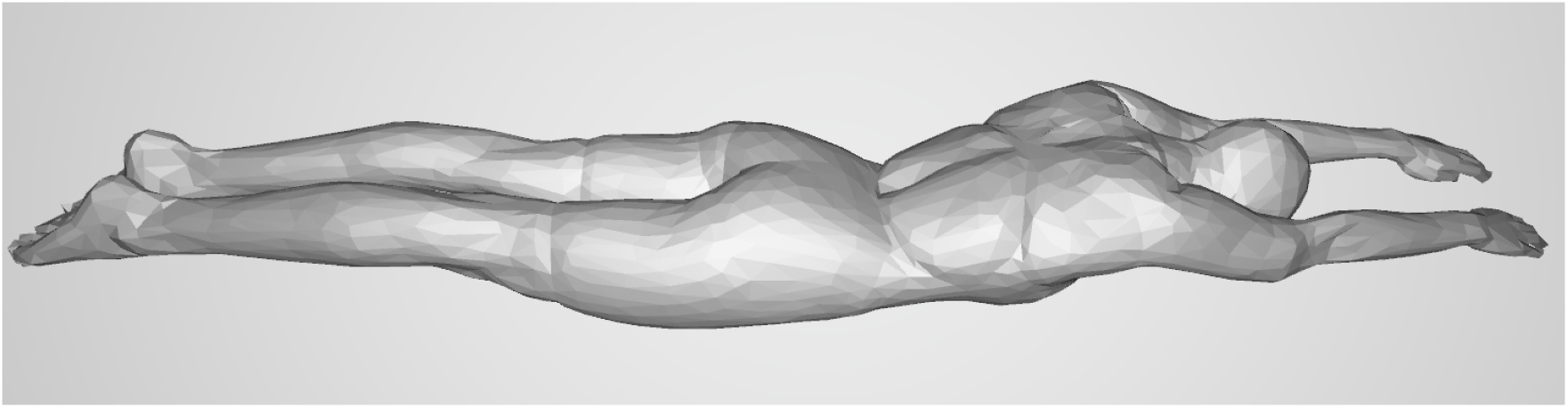
The full geometry of the human female swimmer used in the numerical simulations presented.

The wave resistance of the full geometry is mostly smaller than the truncated geometry, as shown in Figure 11. Figure 11a shows an increase in wave resistance at 1.50 m/s (*Fr* = 0.379) for the cut geometry. The wave resistance of the full geometry is seen to increase between 1.50 − 1.75 m/s (*Fr* = 0.314 − 0.367), Figure 11b. Some relative differences in wave resistance between the full and truncated mannequin geometry can be explained by the similarity of Froude numbers, but differences remain in the shapes of the contours.

**Figure 11:**
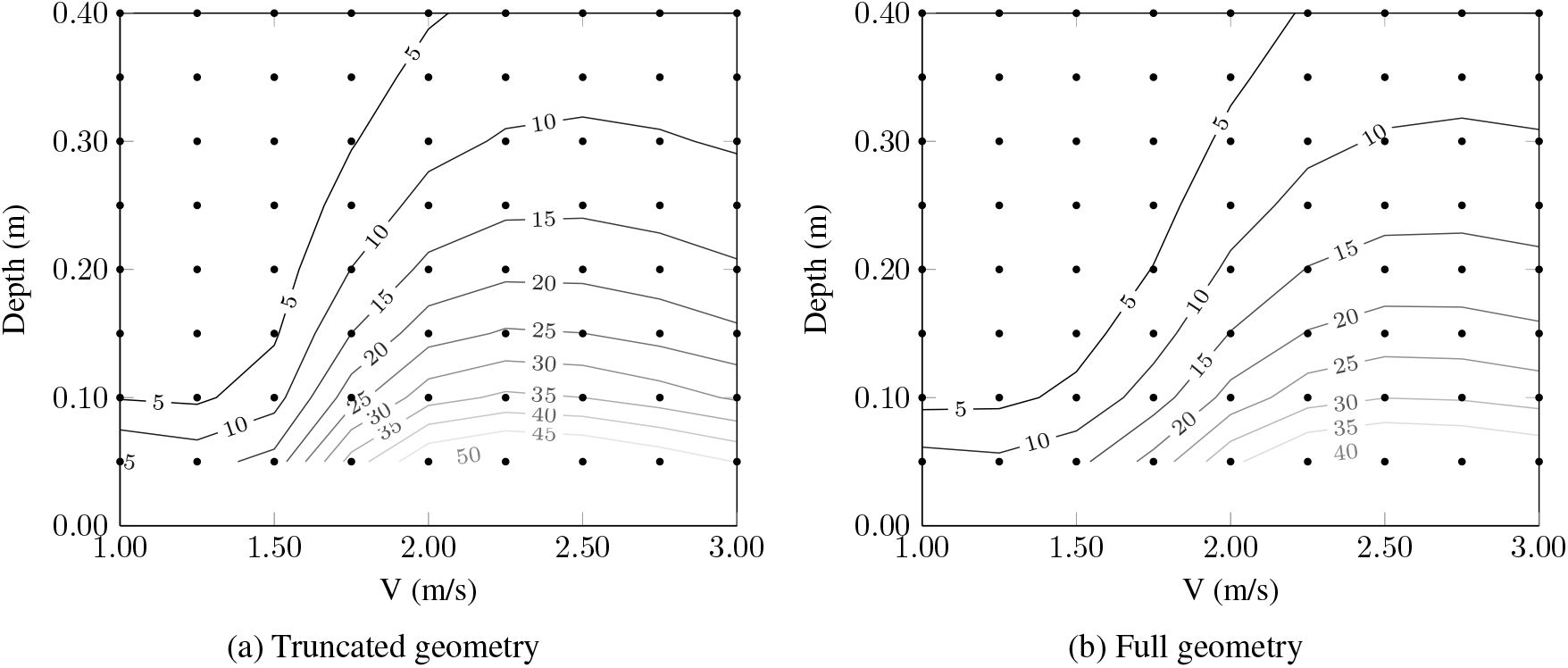
Wave resistance (N) for the a) truncated geometry and 1.00 and b) full geometry. Simulations were conducted at speeds between 1.00m/s and 3.00m/s in increments of 0.25m/s and at depths between 0.05m in increments of 0.05m to 0.50m and up to a depth of 1.00m in increments of 0.10m.

### 5.3 Influence of pool dimensions

Understanding how the channel size influences the wave resistance of a swimmer contextualises experiments and assists athletes with their understanding of how their environment influences their performance. The ability to relate the change in wave pattern to the change in wave resistance is of key interest to coaches, swimmers and academics alike. The wave resistance of the full geometry was calculated for an open Olympic swimming pool and for a single lane within an Olympic swimming pool. The dimensions for the Olympic swimming pool and the single lane are recorded in Table 3.

Figure 12 shows that the wave resistance was increased in the restricted conditions of a single swimming pool lane. The influence of shallower and narrower channels is to increase the magnitude of the wave resistance at relatively earlier speeds and deeper depth. The results in this Figure are the first results which quantify the magnitude of wave resistance that a real athlete would experience in a swimming pool.

**Figure 12:**
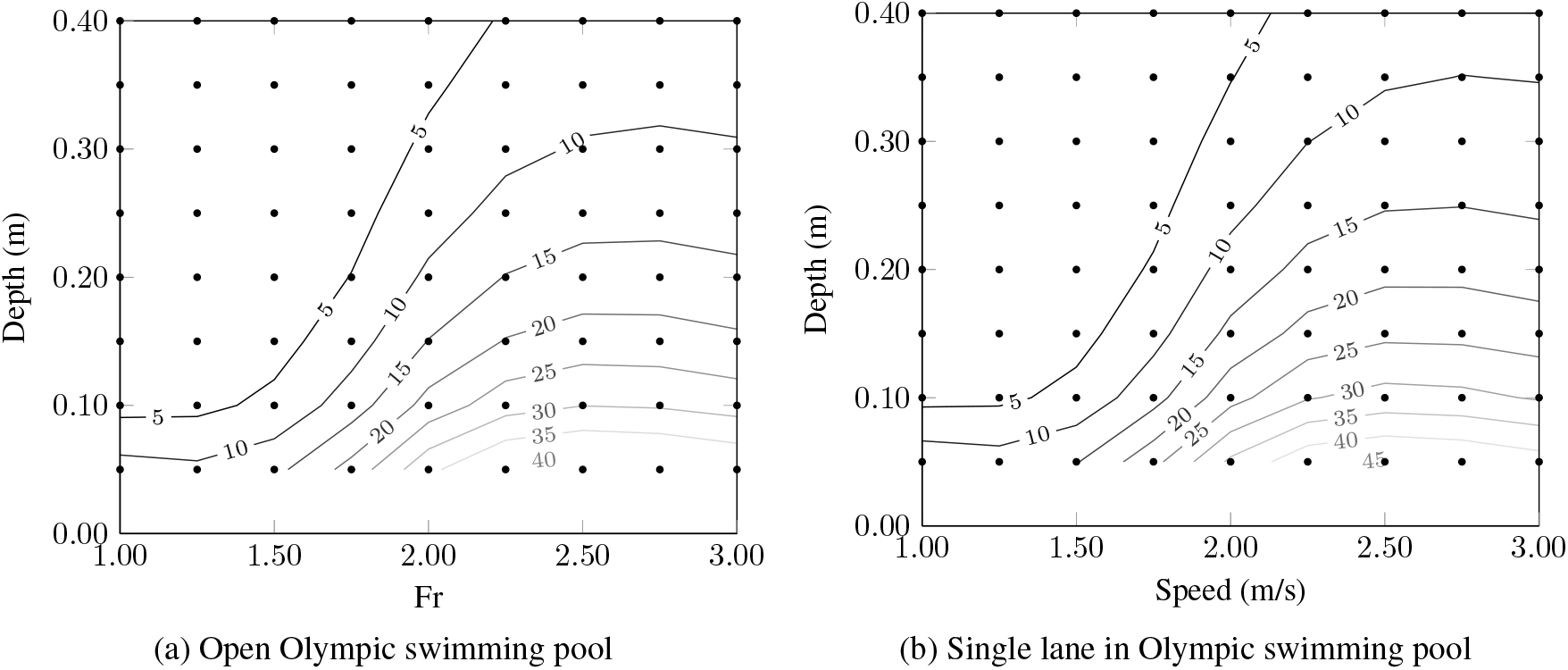
Wave resistance (N) for the full geometry in an open Olympic swimming pool and for a single lane over speeds increasing in increments of 0.25m and depths increasing by 0.05m from *D* = 0.05m to *D* = 0.5m and by 0.1m until a depth of 1.00m.

The wave resistance of the full body geometry in a swimming lane is plotted in Figure 12b. The wave pattern at a depth of 0.05m for a speed of 2.5m/s is shown in Figure 13 and the 0.40m depth is seen in Figure 14. As the immersion of the athlete is increased there is reduction in the size of the divergent wave pattern. The divergent wave pattern at *D* = 0.05m in Figure 13 has disappeared by *D* = 0.40, shown in Figure 14. The overall reduction in magnitude of the wave pattern height and the disappearance of the divergent wave pattern can be associated with the reduction in the magnitude of the wave resistance. The association between the disappearance of the divergent wave pattern and the largest reduction in wave resistance could be a useful cue for coaches to use with athletes poolside.

**Figure 13:**
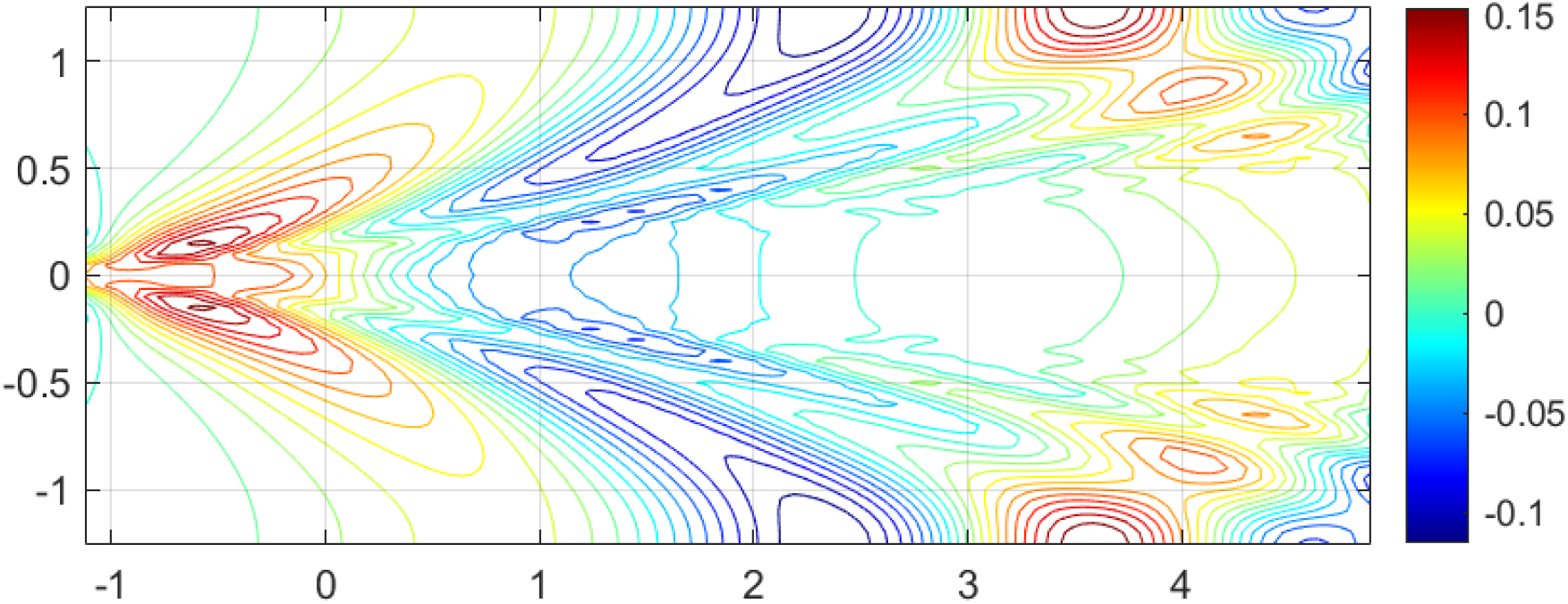
Wave pattern generated at a depth of 0.05m at a speed of 2.5m for the full geometry mannequin in an Olympic swimming pool lane.

**Figure 14:**
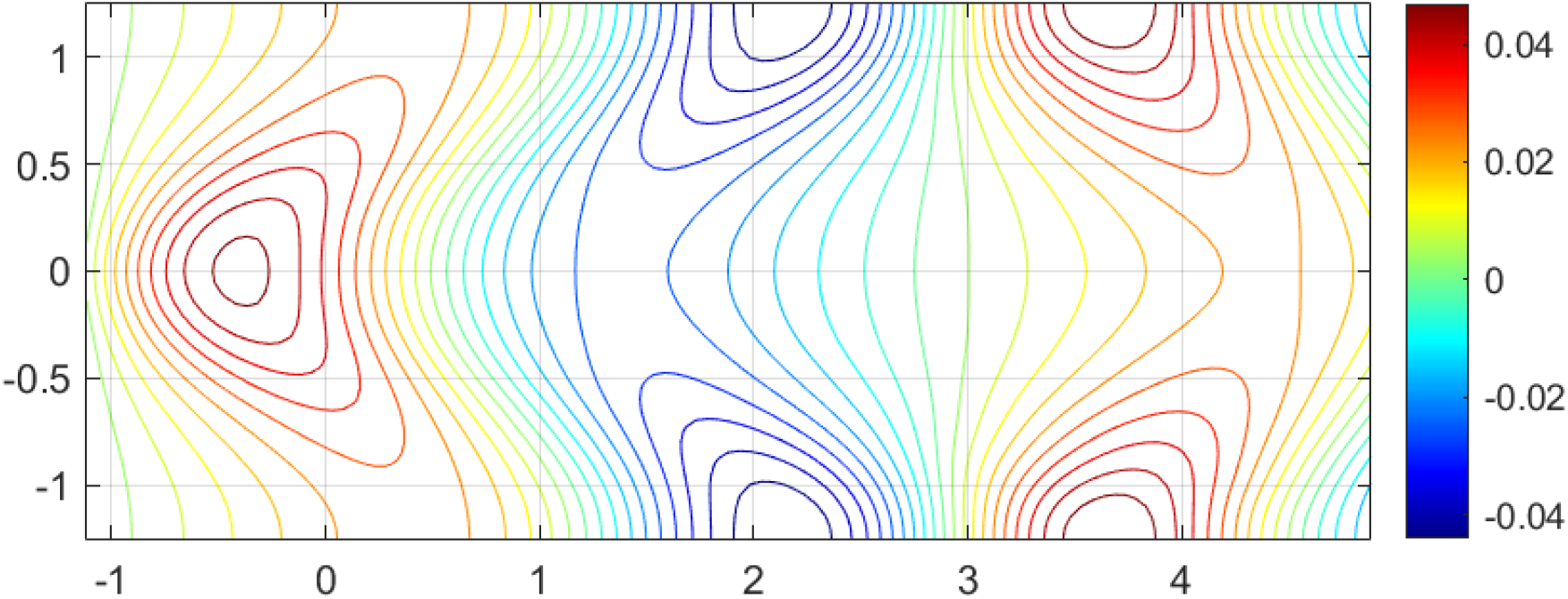
Wave pattern generated at a depth of 0.40m at a speed of 2.5m for the full geometry mannequin in an Olympic swimming pool lane.

Simulating the wave resistance within a single swimmer lane is based on the assumption that there are identical swimmers either side of the lane which are moving at the same speed and depth as the swimmer being modelled. It is likely that this assumption does not hold in competition as some wave energy is lost in the damping of the lane float lines and athletes will not be mirroring each other exactly.

### 5.4 Influence of trim

During competitive swimming athletes often adopt a technique where their body is at an angle to the free surface. Investigating the influence of different trim angles on wave resistance will illustrate the impact of the swimming styles that different athletes adopt. The influence of trim is investigated by rotating the full mannequin geometry about the centre of volume to bring the head or toes closer to the surface. Positive angles describe the clockwise rotation about the y-axis of the swimmer about the centre of volume, bringing the head and shoulders closer to the surface.

Numerical simulations investigated the how varying the trim angle between −15° and 15° in increments of 5° influenced the wave resistance at a depth of 0.40, the depth identified as being the transition between influential and negligible amounts of wave resistance. Figure 15 shows how different trim angles dominate the wave resistance over different speed ranges. There is a very low level of wave resistance recorded at a speed of 1.00m/s. All trim variations from 0° are seen to cause increases of between 1 − 4N above the 0° case, an increase of between 10 − 66%. The results are more complex between 1.50 − 2.50 m/s, where higher trims dominate are relatively larger than the associated lower trim angles. Any deviation from horizontal is seen to cause an increase in the wave resistance at a competition glide speed of 2.50 m/s, indicating that care must be taken in order to maintain a horizontal body position.

**Figure 15:**
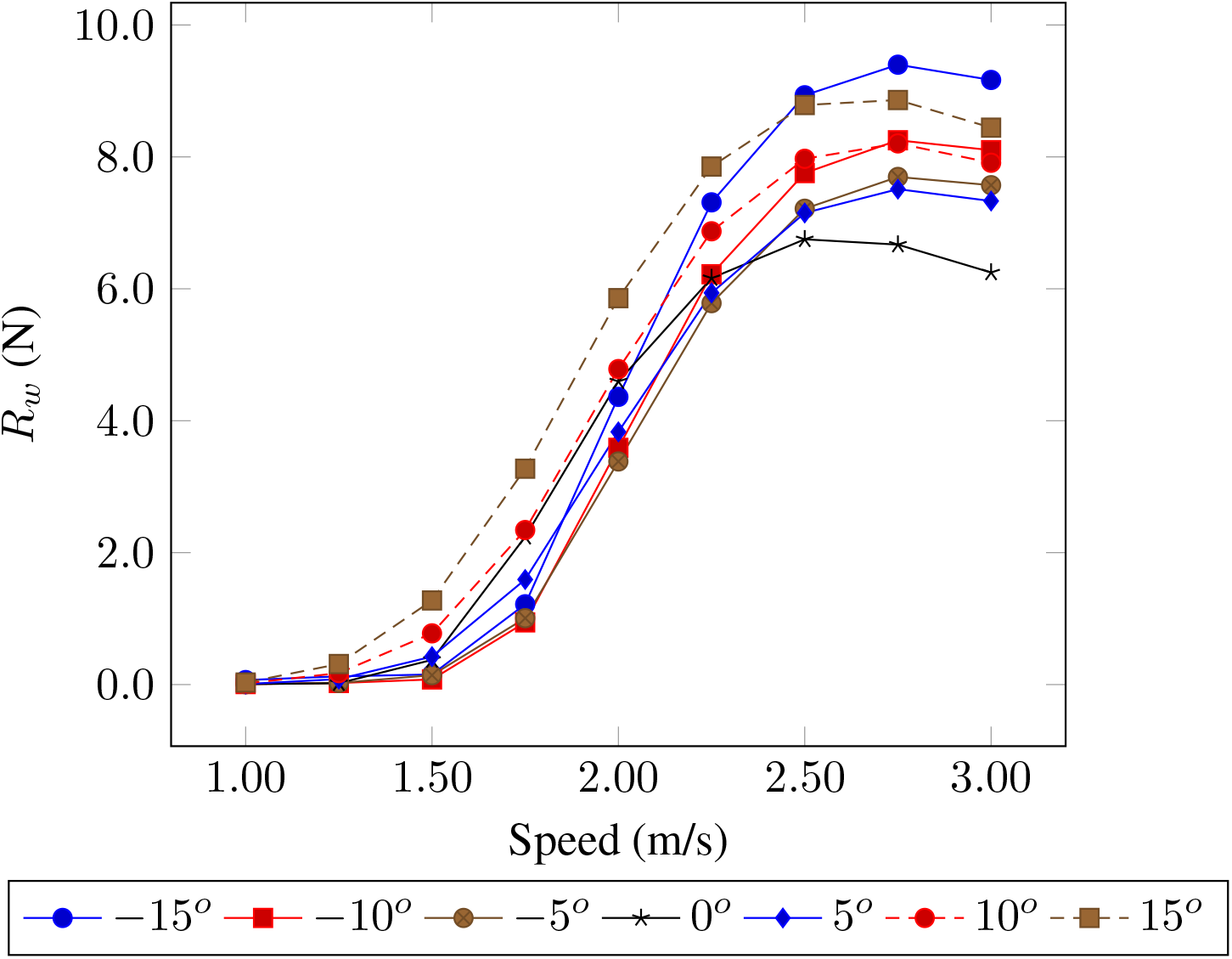
The wave resistance for different trims of the swimmer geometry in an Olympic swimming pool travelling at 2.50m/s at a depth of 0.40m.

## 6 Discussion

A relationship between wave resistance and the immersed depth of a swimmer has been known to exist for several decades. There has been no previously published experimental investigation which has directly quantified the wave resistance component of the resistance of a swimmer as a function of depth. Furthermore, there has been no direct validation of theoretical predictions of wave resistance against experimental results. Validating the use of thin ship theory on swimmer geometries allows problems which were previously computationally intractable to be solved such as quantifying the influence of depth or drafting.

This approach taken in this paper has been to validate the use of TST by comparing the wave resistance in the wave pattern generated by a truncated mannequin towed at a constant speed of 2.5m/s to numerical predictions. A similar trend between the theoretical and experimental wave resistance predictions were found, although there were differences in the distribution of wave resistance across all wave components. The swimmer mannequin contributed to the magnitude of the transverse wave set which diminished at higher depths. No significant reduction in resistance was found to be achieved below a depth of 0.40m. The disappearance of the divergent wave pattern is associated with a reduction in wave resistance.

The previous experimental study into swimmer wave resistance inferred that that the wave resistance was approximately 5% of the total resistance at a depth of 0.5m and a speed of 1m/s (approx 5N) and again for a speed of 2m/s at and immersion of 0.7m [1]. These results broadly agree with the present study which finds that the wave resistance is below 5N in both these situations, based on thin ship theory. The trends seen in Figure 6, [1] have been replicated in this study, although there are some key differences between each study which must be considered.

The mannequin used in the previous experimental study was male and did not necessarily maintain a fixed trim as a function of speed throughout each experiment, the trim attained was not recorded. The flume used has smaller dimensions (2.5 m wide and 1.5 m deep) than the Solent towing tank. The passive drag of a female mannequin has been shown to be larger than a male mannequin [21]. The increase in drag associated with the different mannequin shape and the relative reduction in channel size explain why the additional forces recorded previously are larger [1].

The additional resistance recorded in previous experiments, [1], has been shown to reduce above speeds of 1.8m/s below a peak of 60N at a velocity of 1.6m/s, this contrasts with the present research where the wave resistance is shown to increase with speed. The difference can perhaps be explained by the fixed trim of the mannequin used in this research and the variable trim used previously. Section 5.4 demonstrated that any change in trim over most speeds causes an increase in wave resistance relative to a fixed trim. The trim of an athlete or mannequin must be reported in order to compare results.

A numerical investigation using linear, slender body theory quantified the wave resistance generated by single and multiple swimmer like forms [14]. It was found that whilst the swimmer was near the free surface (*h* = 0.00 − 0.20m) there was significant wave drag which reduced to a negligible amount beyond *h* > 0.40m. The experimental results presented in the present research broadly agree with these results, however there are differences due to the mannequin geometry. The geometry used in [14] has no legs, arms or realistic body detail. Therefore, the idealised swimmer shape has a different Froude number, hence wave pattern, relative to the realistic swimmer geometry used in this research.

The mannequin used in the present research is rigidly fixed to the towing carriage support structure. The use of a mannequin reduces the number of error sources and inconsistencies found with running experiments on human subjects. These errors include the ability to maintain a steady depth, trim and lateral position of a swimmer throughout an experiment. It is possible to estimate the drag of a swimmer using a mannequin to provide a realistic body shape which reduces the sources of error associated with human athletes, which have been encountered in previous research [13].

The wave probes were set up to capture the longest trace of wave height possible, however there are issues with the reflection of the waves on the tank wall, the size of the tank and the calibration of the wave probes. The position of the beaches mean that energy is lost in the wave system as it is reflected from the wall, therefore the wave traces have to be cropped before the reflected waves pass through the probes. If the tank was longer and wider it would be possible to record a longer wave pattern and increase the accuracy of the analysis. Improvements in the experimental accuracy can be obtained through conducting experiments in a larger tank with flat sides.

The error associated with the wave probe calibration is quantified through quantifying the change in experimental wave resistance as a consequence of altering the calibration factor. The conversion factor was varied in fixed increments of −1, −0.1, 0.1, 1 mm/V and was used to estimate the wave resistance for the wave cuts at immersed mannequin depths of 0.05m and 0.4m using the matrix method and an individual wave probe.

The changes in calibration factor considered are associated with a small standard deviation in the wave resistance for both depths and wave probe combinations, Table 5. There is a larger standard deviation associated with the wave resistance calculated using a single wave probe relative to the matrix method. The change in calibration factor alters the magnitude of the wave resistance recorded but does not alter the trend. It can be seen that the results change by less than 5% for the largest changes in calibration factor, Table 5. These results indicate that there would have to be high levels of uncertainty associated with the calibration factor in order to alter the results, and no change will be seen in the general trend.

**Table 5:**
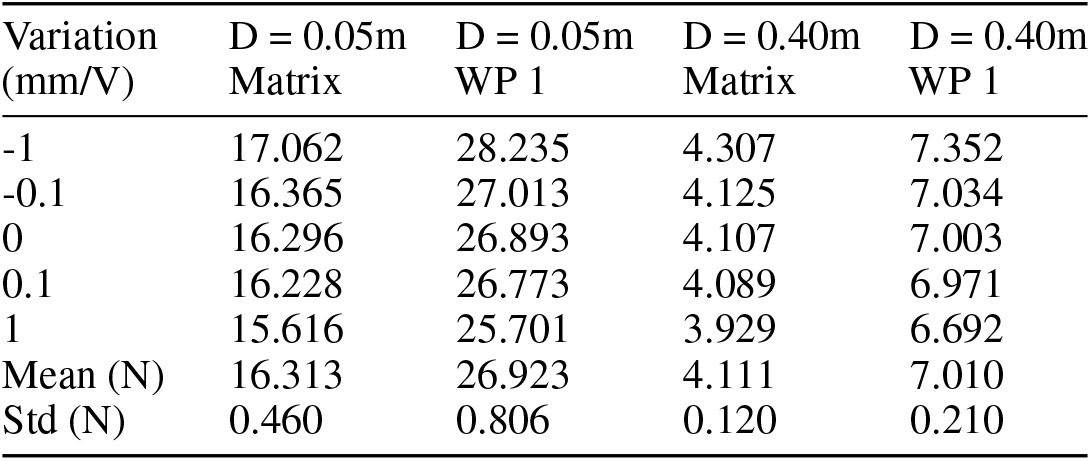
Influence of variation in calibration factor on the estimated wave resistance from a single and multiple wave probes.

| Variation<br>(mm/V) | D = 0.05m<br>Matrix | D = 0.05m<br>WP 1 | D = 0.40m<br>Matrix | D = 0.40m<br>WP 1 |
| --- | --- | --- | --- | --- |
| -1 | 17.062 | 28.235 | 4.307 | 7.352 |
| -0.1 | 16.365 | 27.013 | 4.125 | 7.034 |
| 0 | 16.296 | 26.893 | 4.107 | 7.003 |
| 0.1 | 16.228 | 26.773 | 4.089 | 6.971 |
| 1 | 15.616 | 25.701 | 3.929 | 6.692 |
| Mean (N) | 16.313 | 26.923 | 4.111 | 7.010 |
| Std (N) | 0.460 | 0.806 | 0.120 | 0.210 |

## 7 Conclusions

This paper has quantified how the wave resistance of a swimmer reduces as the depth increases. The validation of thin ship theory has been achieved by comparing the experimental and numerical wave resistance generated by a swimmer at varying depths. Small discrepancies between experimental and theoretical predictions were found to originate from the inability to capture the energy lost in the breaking waves in the divergent wave system. The wave resistance reduces significantly below a depth of 0.40 for competition swimming speeds. The disappearance of the divergent wave system from the wave pattern generated by the athlete could be a useful cue for athletes and coaches seeking to optimise their performance.

The contribution of different body parts to the wave pattern was studied by varying the trim of the full swimmer geometry and comparing the wave resistance at a level trim to the truncated swimmer geometry. Altering the geometry of the swimmer by removing the legs increases the wave resistance between speeds of *Fr* = 0.30 − 0.50. At *Fr* < 0.30 and *Fr* > 0.60 the wave resistance for both geometries appear to follow similar trends. It appears that the head and shoulders create a more influential wave pattern below speeds of 2.00m/s and that any deviation from a level trim increases the wave resistance at higher speeds.

Development in this research would include experimental measurement of the resistance of real athletes at various depths in a swimming pool. Measuring the resistance of athletes at various depths would quantify how realistic the use of a mannequin is. It would also be possible to investigate how different athlete proportions contribute to the total resistance and how this impacts wave resistance. Evaluating the influence of different mannequin geometries such as male or female and varying length/volume ratios will also identify how different athletes may optimise their technique for competition. Furthermore, quantifying the side and lift forces acting on the mannequin will improve the understanding of the forces acting on a submerged swimmer.

## Supporting information

Supporting data and analysis programs

## 8 Ethics

Identifiable human features are not recorded in the photos therefore ethics review and approval was not required.

## 9 Data accessibility

Data from the experiments are included in the electronic supplementary material. The Matlab packages used to analyse the data are also included. The 3D geometric model of the mannequin is not released as it has identifiable features.

## 10 Authors acknowledgements

T.D. collected data, performed numerical and experimental analysis and drafted the manuscript. T.D, D.T., J.B., D.H., S.T. conceived, designed and coordinated the study and helped draft the manuscript. D.T. built the mathematical model.

## 13 Acknowledgements

We want to acknowledge Christopher Phillips and Apostolos Grammatikopoulos with their assistance in designing and manufacturing the experimental equipment. We also thank Dorian Audot, Isobel Thomas and Christopher Phillips with their assistance with experimental data suggestion. Apostolos Grammatikopoulos contributed edits and

