## Supporting data and analysis programs for "Quantifying the wave resistance of a swimmer": Calibration.m

```
clear all;
close all;
clc;

[file path]=uigetfile('*.mat');

% Calibration.m
% 
% File to allow the calibration of wave probes.
% Click "run" to access the GUI to calibrate the user interface.
% The user will then click the voltage levels that respond to the following changes in displacement;
% 
% 1. -50 mm
% 2. 0 mm
% 3. 50 mm
% 
% At this point a graph of voltage vs displacement will be plotted and the "rate" will be returned in the command line.
% The channel for measurements can be changed immediately below this
% comment. The number corresponding to each channel is displayed below.
% 2 = WP1, 3 = WP2, 4 = WP3, 5 = WP4

%%%%%%%%%%%%%%%%%%%%%%%%%%%%%%%%%%%%%%%%%%%%%%%%%%%%%%%%%%%%%%%%%%%%%%%%%%%%%
%%%%%%%%%%%%%%%%%%%%%%%%%%%%%%%%%%%%%%%%%%%%%%%%%%%%%%%%%%%%%%%%%%%%%%%%%%%%%
channelI = 4; % change number dependent on what channel is required
%%%%%%%%%%%%%%%%%%%%%%%%%%%%%%%%%%%%%%%%%%%%%%%%%%%%%%%%%%%%%%%%%%%%%%%%%%%%%
%%%%%%%%%%%%%%%%%%%%%%%%%%%%%%%%%%%%%%%%%%%%%%%%%%%%%%%%%%%%%%%%%%%%%%%%%%%%%
%DELIMITER = '\t';
%HEADERLINES = 25;
% Import the file
%S = importdata([path file], DELIMITER, HEADERLINES);
S = load([path file]);
A=S.data;%unfiltered data
sr=100;%sample rate
time=0:1/100:(length(A)-1)/100;

%% lowpass Filter
[b,a]=butter(2,2/(sr/2),'low');
AF=filtfilt(b,a,A);%filtered data

%% Get calibration rate
figure;
plot(AF(:,channelI)) 
data=ginput(3);
close

data(:,1) = [-50 0 50];

WaveEl = data(:,1);
Volt = data(:,2);

figure;
scatter(Volt, WaveEl);
p = polyfit(Volt, WaveEl , 1);
rate = p(1)
hold on;
plot(Volt, WaveEl,'r-')
```
