## Supporting data and analysis programs for "Quantifying the wave resistance of a swimmer": GET_waveCutForWaveSis.m

```
% GET_waveCutForWaveSis
% 
% File to produce a wave cut from experimental data recorded using wave probes.
% User must input the "rates" measured from a calibration procedure. 

 
[file path]=uigetfile('*.mat');
S = load([path file])
A=S.data; %unfiltered data
sr=100; %sample rate
time=0:1/100:(length(A)-1)/100;

%% lowpass Filter
[b,a]=butter(2,2/(sr/2),'low');
AF=filtfilt(b,a,A);%filtered data

    
%% Calibrate - UPDATE RATE
probe_1_rate = -50.6066;
probe_2_rate = -47.7376;
probe_3_rate = -44.6288;
zero=mean(AF(1:500,:));
% zero=mean(A(1:500,:));
wave1=(A(:,2)-zero(2)).*probe_1_rate*0.001; %Furthest away from model
wave2=(A(:,3)-zero(3)).*probe_2_rate*0.001; %middle
wave3=(A(:,4)-zero(4)).*probe_3_rate*0.001; %Nearest the model?
wave4=(A(:,5)-zero(5)).*0*0.001; %THIS Probe is not being used!!! ( It's the one nearest to the mpode


%WAVECUT.WAVE=[wave1 wave2 wave3 wave4];
WAVECUT.WAVE=[wave3 wave2 wave1]; % original probe order - don't fiddle with this as I (or you) don't understand how this is loaded
%WAVECUT.WAVE=[wave1 wave2 wave3]; % wave probes in the closest to model first order (this was put in reverse previously)
WAVECUT.WIDTH=3.7; %Tank width
WAVECUT.DEPTH=1.8; % Tank depth 
WAVECUT.L=1.6; %NPL = 1.6m QM = 2.48m
WAVECUT.WSA=1.7063; %wetted surface area (m2)
WAVECUT.U=2.527; % measured speed (m/s)
WAVECUT.N=3; %Number of probes
WAVECUT.Y=[1.11 1.207 1.291]; % distance from centerline - closer to model first
WAVECUT.SR=100; %Sampling Rate
%WAVECUT.RT=results(i,3);
   
%    Save WAVECUT
save([path file,'WAVECUT','.mat'],'WAVECUT','-mat');
```
